# Assessing the influence of the Mediterranean diet on dental calculus microbiome composition: a pilot study

**DOI:** 10.1101/2022.06.01.494314

**Authors:** Gabriel Innocenti, Maria Elena Martino, Edoardo Stellini, Adolfo di Fiore, Andrea Quagliariello

## Abstract

Human oral microbiome harbours hundreds of bacterial species, which, during evolution, assembled in different niches, such as the microbiome of saliva, tongue, hard palate, gingiva and dental plaque. Among these, saliva has been the most studied ecological niche of the oral microbiome. In this frame, scientific research has demonstrated that several factors concur to determine the saliva microbial composition, including diet, systemic diseases and drugs. However, the influence of such variables on other oral niches, such as the dental calculus, a calcified derivative of dental plaque, is still unexplored.

During the last decade, dental calculus has become the primary target for evolutionary microbiome studies because of its ability to keep both bacterial DNA and dietary molecules (i.e., proteins and starch). As a consequence, dental calculus is now considered a powerful tool for obtaining information about oral microbiome composition, diet and individual health status in past populations, making it also appealing for clinical research.

Here, we provide the first clinical study, to our knowledge, assessing the inter-variability of dental calculus microbiome in 40 patients. Samples were collected during routine dental inspection and analysed through 16S amplicon sequencing. In addition, we investigated the relationship between oral microbiome and host. Specifically, we tested the influence of Mediterranean diet style on the composition and functional pathways of oral bacteria, with the aim to test the dental calculus as a clinical bio-marker for diet impact and health status.

## Introduction

The interaction between humans and their microbiomes has become one of the main growing interests in human health in the last few decades^1^. Extensive research in different scientific fields, spanning from ecology to evolution and biomedicine, has investigated how the microbiota composition affects human health, and vice versa^2^. In addition, the recent breakthrough in personalized medicine has made it possible to integrate additional data beyond host genetics or microbiome composition, such as human lifestyle data (i.e., daily habits, diet, drug use, and sport activity)^3,4^. Diet is known to be one of the main factors associated with microbiome compositional shifts, explaining 60% of such variations^5^. Furthermore, adherence to a specific diet seems to promote the opportunistic pathogenesis of some microorganisms^6^. In this context, the oral tract microbiome is gaining attention as a new target of omics studies, as it can capture humans’ dietary habits, immune responses, and environmental factors^7^. The oral cavity represents one of the richest substrates for microorganisms to grow and establish complex microbial communities^8^. Several studies have investigated how shifts in the oral microbiome composition correlate with a higher risk of diseases such as periodontitis, caries, and even bacteremia events^9^, in which diet clearly plays a key role. Previous research comparing changes in microbial communities driven by Mediterranean, western, or vegan diets has focused almost exclusively on gut microbiota^10,11^. As a consequence, the influence of dietary regimen on the human oral microbiome remains largely elusive.

Recently, molecular anthropology and archaeology have greatly improved our knowledge regarding the evolution of human–microbe symbiosis during human history^12,13^. The ability to retrieve conserved DNA from ancient samples has resulted from the analysis of tooth-associated dental calculus, a mineralized derivation of the commonly known dental plaque. Dental calculus has been demonstrated to be an extremely rich resource, harboring a variety of biological molecules (i.e. host and microbial DNA, proteins, starch grains, and metabolic compounds^14^), which has allowed us to obtain important information regarding the dietary habits and health status of populations from the past. In addition, as recently observed, dental calculus has overcome many of the problems normally encountered during ancient DNA conservation in other tissues^15^, becoming the most common source of ancient microbiomes. However, to our knowledge, dental calculus has rarely been considered in clinical studies on modern microbiota. Previously, a model of plaque formation was proposed to explain the establishment of the oral microbiome community, identifying the succession of early and late colonizers^16–20^. Recently, new evidence seems to reject the proposed model in favor of a more complex structure and composition in which species traditionally belonging to both early and later colonizers are mixed together in large aggregates; once bound to the teeth, this mixed microbial community starts biofilm formation, which then engulfs the bacterial single static cells found in saliva^21^. However, to date, we still lack information regarding the possible microbial biodiversity harbored in such a specific ecological niche as the dental calculus, and regarding whether it presents a certain level of inter-individual variability or, conversely, whether the dental calculus ecological niche flattens microbial biodiversity. Moreover, we currently have no evidence to disclose whether lifestyle and health variables are able to shape its composition, as observed in microbiomes found in other body niches. In this study, we analyzed the microbiome composition of modern dental calculus in order to define (i) the level of inter-individual variability and (ii) the role of the diet in driving its structure. To this end, taxonomic diversity and microbial community functional profiles were investigated by applying 16S amplicon sequencing methodology to 40 omnivorous female and male subjects, ranging from 20 to 77 years of age, following a Western lifestyle. This work can be considered a pilot study testing how the dental calculus microbiome could be exploited to obtain relevant clinical information on diet effects and patient health status. In addition, it provides a modern reference for human microbiota evolutionary studies.

## Materials and methods

### Sample collection and surveys

Dental calculus samples were collected after Ethics Committee approval (Prot. N°AOP2258) at the Unit of Dentistry of the Hospital of Padova (Italy) during routine inspections. Patients were informed about the study aims and provided their informed consent. A questionnaire was submitted to each patient, requesting anthropometric, lifestyle, health status, and dietary information. A published dietary survey was applied in order to collect specific data about subject diet and adherence to the Mediterranean diet^22^. Samples were collected in 1.5ml Eppendorf tubes and directly transferred to the biomolecular laboratory at the Department of Comparative Biomedicine and Food Science of the University of Padova (Italy).

### DNA extraction and sequencing

Genomic DNA was extracted using a Qiagen PowerSoil kit, following the manufacturer’s instructions (Qiagen, Hilden, Germany). In order to maximize the DNA extraction yield, the dental calculus was fragmented using a sterile micro-pestle. The amount of extracted DNA was estimated through Qubit 4 Fluorometric Quantification using the Qubit™ dsDNA High Sensitivity Assay (Thermo Fisher, USA). Samples were then sent to the BMR-Genomics s.r.l. facility (Italy) for 16S rRNA sequencing of the V3-V4 regions, using primers 341F (5-TCGTCGGCAGCGTCAGATGTGTATAAGAGACAGCCTACGGGNBGCASCAG-3’) and 805R (5’-GTCTCGTGGGCTCGGAGATGTGTATAAGAGACAGGACTACNVGGGTATCTAATCC-3’) from Takahashi et al. (2014)^23^, using Taq Platinum HiFi (Thermofisher, USA). The cycling conditions consisted of an initial template denaturation at 94°C for 1 min, followed by 25 cycles of denaturation at 94°C for 30s, annealing at 55°C for 30s, and extension at 68°C for 45s, followed by a final extension at 68°C for 7min.

PCR amplicons were purified using Thermolable Exonuclease I (NEB), diluted at a ratio of 1:2 and amplified following the Nextera XT Index protocol (Illumina, Inc.). The amplicons were normalized using a SequalPrep™ Normalization Plate Kit (Thermo Fisher Scientific Inc.) and multiplexed.

The pool was purified with Agencourt XP 1X magnetic beads (Beckman Coulter, Inc., CA, U.S.A.), loaded on the MiSeq System (Illumina, Inc., USA), and sequenced following the V3 - 300PE strategy. Raw reads were deposited in the BioProject database under accession number PRJNA828039.

### Read processing and taxonomic analysis

The produced sequences were analyzed according to Callahan et al. (2016)^24^. Paired-end sequenced reads were processed by quality filtering using the package *dada2* (v. 1.14.1) on R version 3.6.3. Reads were trimmed according to their base quality at the 5’ ends (where the average quality decreases), using a size of 300bp for the forward and 270bp for the reverse. Then, the fragments were de-replicated, resulting in the collapse of every identical sequence into a unique one as a representative, and finally, they were merged together. The resulting sequences were then clustered using the Amplicon Sequence Variants approach (ASV), removing chimera sequences at the end of the process. ASVs were assigned to their respective taxonomical classification until genus level using a training set provided by the SILVA 16S Database (NR98 version 138.1). Another filtering strategy was applied by removing all taxa for which the prevalence was under the threshold of 0.05% (out of 1) for the entire dataset, resulting in the exclusion of all the unique sequences (singletons). Taxon raw counts (ASVs) were transformed into relative abundance levels, reporting the percentage of each one with respect to the total number of sequences accounted for in every sample. Taxonomic information, ASV counts, and sample metadata were merged together using the *phyloseq* package (v. 1.30.0), and a set of explorative analyses was performed to investigate the taxonomical abundances for each individual and sample’s inter- and intra-variability regarding microbiome composition.

The core microbiota was determined using the *microbiome* package (v 1.8.0), using a 0.1% compositional abundance threshold and a prevalence of 0.5, focusing on the genus level. A principal coordinate analysis (PCoA) using the Bray–Curtis distance was used to observe the main differences in our dataset. Then, we performed a cluster analysis based on the taxonomic differences among samples. To do so, a hierarchical clustering algorithm was applied to the whole dataset, and the optimal number of clusters was computed using the gap statistic, applying the PAM clustering algorithm, with a maximum number of 20 clusters and 500 bootstraps. Moreover, we evaluated the goodness of the gap statistic using the *clusGap* function from the *cluster* (v. 2.1.3) package in R. An ANOVA was additionally executed on the resulting clusters using the Shannon and observed indexes. In order to understand which taxa significantly changed in terms of abundance and diversity along the resulting clusters, a Kruskal–Wallis test was also applied to the whole genus-level community. Finally, computing the LDA effect size with LEfSe^25^ allowed us to assign each of the significant taxa to the cluster for which they were representative.

### Diet metadata integration

Food intake associations with the host microbiome were checked using all available metadata on daily consumption of meat, vegetables, fruit, alcohol, olive oil, milk and cheese, cereals, legumes, and fish. In addition, the Mediterranean score of each sample was considered as part of the microbiome correlation analysis, as done previously^26^. Daily consumption values, represented by nominal and qualitative variables in our dataset, were then transformed into ordinal values, allowing a quantitative evaluation: we assigned ‘<1’ to 0, ‘1–2’ to 1, and ‘>2’ to 2. The association between diet type and microbiome composition was checked through a combined statistics approach. First, based on the detrended correspondence analysis (DCA) result, redundancy analysis (RDA) was found to be the most appropriate method with which to investigate the relationship between dietary and taxonomic variables. A permutation analysis to assess the significance of dietary variables that fitted with the ordination was applied through the *envifit* function from the *vegan* (v. 2.4-2) package in R. Then, the Mediterranean scores of each cluster were analyzed using an all-versus-all t-test to find possible associations between different diet styles and microbial communities. Finally, a Spearman test was performed to determine which and to what extent each genus-level taxon correlated with Mediterranean scores.

### Functional analysis

To assess the gene content and functional profiles of each sample’s microbiome, Picrust2 softwar^e27^ was applied to the pre-filtered ASV sequences. Gene families and relative abundances were computed using the KEGG Ontology database^28^, enabling the collapse of these functions to the pathway levels 2 and 3 (L2, L3) based on the KEGG mapping tree. The normalized counts for each function were analyzed as performed for the taxonomic data; hierarchical clustering was used to detect the main differences in the functional profiles of each microbiome using the Bray–Curtis distance, and a gap statistic was obtained using the k-means clustering algorithm to predict the optimal number of clusters. Then, a LEFSe test was performed to determine the significant changes in pathways for both L2 and L3 functions between the obtained groups. Finally, the morphologies of the function-based tree and the taxonomy-based tree were compared using the ‘tanglegram’ function in the *dendextend* package v1.15.2, with the aim to understand which taxonomic clusters fell into diverse functional groups and to derive the relations between microbiome activity, microbial composition, and sample dietary information.

## Results

### Patient metadata reveal a diverse range of food consumption

Our dataset was composed of 40 samples comprising 13 male and 27 female participants of varying ages, ranging from 20 to 77 years old, with an overall mean of 48.1 (male mean age = 47.69, female mean age = 48.34) and a median of 50. The age distributions of males and females were similar to ensure that every age range and sex was well represented (Figure 1A). The majority of patients (39 out 40) were Italians, mainly from Northern Italian regions (37 out of 40, corresponding to 92.5%), with 2 of them originating from South Italy (Sicily), while the remaining participant was Spanish (Table 1).

**Figure 1.**
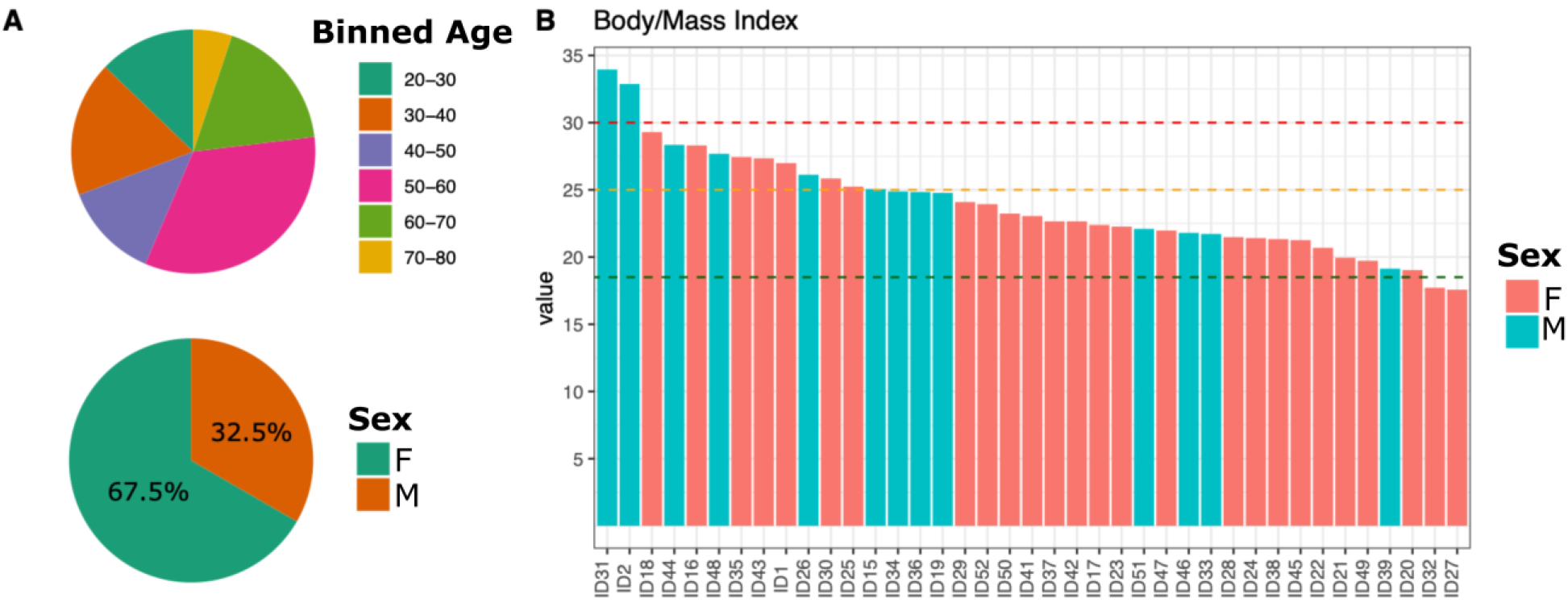
**A.** Pie charts showing the dataset composition in terms of age and sex variables. Age was binned using 10-year ranges. **B.** Barplot showing the body mass index distribution in the cohort. Dashed lines indicate BMI ranges according to WHO Europe: the green line indicates the lower limit for a normal-weight BMI, while the orange line represents the upper limit of the same; the red line indicates the limit between overweight and concrete obesity.

Anthropometric information (e.g., height and weight) was collected through a questionnaire for each patient and used to obtain the body mass index (Table 1). The mean for the overall values of BMI was 23.85, which falls in the normal weight score range according to the World Health Organization (WHO) Europe standard (https://www.euro.who.int/en/). Specifically, the mean BMI of males was 25.63, which is considered slightly above the pre-obesity threshold of 25, while for females, the average BMI was 22.95, corresponding to a normal weight. Within the dataset, only two males (ID2 and ID31) showed obesity conditions, surpassing the value of 30 in the BMI index scale, and also recognizing their obesity in the survey (Table 1 and Figure 1B). In particular, ID2 also showed complex clinical reports, suffering from glaucoma and diabetes. There were also two cases of underweight (ID27 and ID32), both in females, presenting a BMI score lower than 19 (Table 1 and Figure 1B). Despite these specific cases, no significant difference was detected in sex distribution within the binned age groups and BMI (chi-squared test p>0.05).

Data regarding dietary habits showed a very homogenous profile, considering that 39 out 40 individuals were omnivores, but still presented a diverse range of food preferences and daily consumption (Supplementary Table S1). For this purpose, the Mediterranean score assignment strategy was used to define the level of each individual’s closeness to the Mediterranean diet, showing different diet styles (Supplementary Figure S1).

**Table 1.** Information collected from the cohort taking part in the present study: sample age, gender, weight (kg), height (cm), body mass index (BMI), place of birth, and geographical location. Further data are reported in Supp. Table 1.

| Sample-ID | Age | Sex | Weight (cm) | Height (cm) | BMI | Location |
| --- | --- | --- | --- | --- | --- | --- |
| ID1 | 77 | f | 78 | 170 | 27 | North Italy |
| ID2 | 67 | m | 95 | 170 | 33 | North Italy |
| ID3 | 63 | m | 66 | 170 | 23 | North Italy |
| ID15 | 39 | m | 83 | 182 | 25 | North Italy |
| ID16 | 50 | f | 68 | 155 | 28 | North Italy |
| ID17 | 39 | f | 67 | 173 | 22 | North Italy |
| ID18 | 53 | f | 75 | 160 | 29 | North Italy |
| ID19 | 50 | m | 75 | 174 | 25 | North Italy |
| ID20 | 34 | f | 55 | 170 | 19 | North Italy |
| ID21 | 44 | f | 53 | 163 | 20 | North Italy |
| ID22 | 50 | f | 57 | 166 | 21 | Spain |
| ID23 | 58 | f | 57 | 160 | 22 | North Italy |
| ID24 | 34 | f | 59 | 166 | 21 | North Italy |
| ID25 | 54 | f | 63 | 158 | 25 | North Italy |
| ID26 | 21 | m | 80 | 175 | 26 | North Italy |
| ID27 | 25 | f | 49 | 167 | 18 | North Italy |
| ID28 | 29 | f | 55 | 160 | 21 | North Italy |
| ID29 | 57 | f | 68 | 168 | 24 | North Italy |
| ID30 | 45 | f | 67 | 161 | 26 | North Italy |
| ID31 | 62 | m | 110 | 180 | 34 | North Italy |
| ID32 | 60 | f | 50 | 168 | 18 | North Italy |
| ID33 | 63 | m | 65 | 173 | 22 | North Italy |
| ID34 | 38 | m | 87 | 187 | 25 | North Italy |
| ID35 | 39 | f | 72 | 162 | 27 | North Italy |
| ID36 | 50 | m | 85 | 185 | 25 | North Italy |
| ID37 | 56 | f | 58 | 160 | 23 | North Italy |
| ID38 | 62 | f | 48 | 150 | 21 | North Italy |
| ID39 | 20 | m | 62 | 180 | 19 | North Italy |
| ID40 | 29 | f | 56 | 175 | 18 | North Italy |
| ID41 | 62 | f | 62 | 164 | 23 | North Italy |
| ID42 | 46 | f | 58 | 160 | 23 | North Italy |
| ID43 | 54 | f | 70 | 160 | 27 | North Italy |
| ID44 | 70 | m | 80 | 168 | 28 | North Italy |
| ID45 | 45 | f | 60 | 168 | 21 | North Italy |
| ID46 | 63 | m | 63 | 170 | 22 | North Italy |
| ID47 | 54 | f | 65 | 172 | 22 | North Italy |
| ID48 | 40 | m | 80 | 170 | 28 | South Italy |
| ID49 | 28 | f | 57 | 170 | 20 | South Italy |
| ID50 | 50 | f | 58 | 158 | 23 | North Italy |
| ID51 | 37 | m | 70 | 178 | 22 | North Italy |
| ID52 | 52 | f | 59 | 157 | 24 | North Italy |

### Dental calculus microbiome composition and diversity

A total of 39 samples were selected for the microbiome composition analysis. One sample (ID40) was discarded because of a poor read count, probably due to the low quantity of extracted DNA (Supplementary Table S2). Focusing on the relative abundance of each taxon, we found 10 phyla, among which Firmicutes was found to be dominant in ~75% of the samples, followed by Actinobacteria, Bacteroidetes, and Proteobacteria (Figure 2A). Descending to genus-level taxonomy, we observed a more complex composition; in particular, the dental calculus microbiome of the whole dataset was populated by a total of 39 genera, mainly represented by *Streptococcus, Tannerella, Fusobacterium, Freticobacterium, Porphyromonas, Bacteroidetes oral taxon sp*., and *Prevotella* (Figure 2B). The core microbiota comprised 11 taxa, including *Streptococcus, Rothia, Fusobacterium*, *Campylobacter*, *Tannerella*, *Treponema*, *Porphyromonas*, and *Actinomyces*, among others (Figure 2C). In particular, *Streptococcus* showed the highest prevalence in our dataset. This is likely due to the high number of species ascribed to this genus that compose the human oral microbiome. Furthermore, the core microbiota showed the presence of all three taxa comprising the red complex, a group of bacteria formed by *Treponema*, *Tannerella*, and *Porphyromonas*. At the same time, the accessory microbiome comprised bacteria commonly found in the oral cavity, such as *Freticobacterium*, *Abiotrophia*, and *Aggregatibacter*.

**Figure 2.**
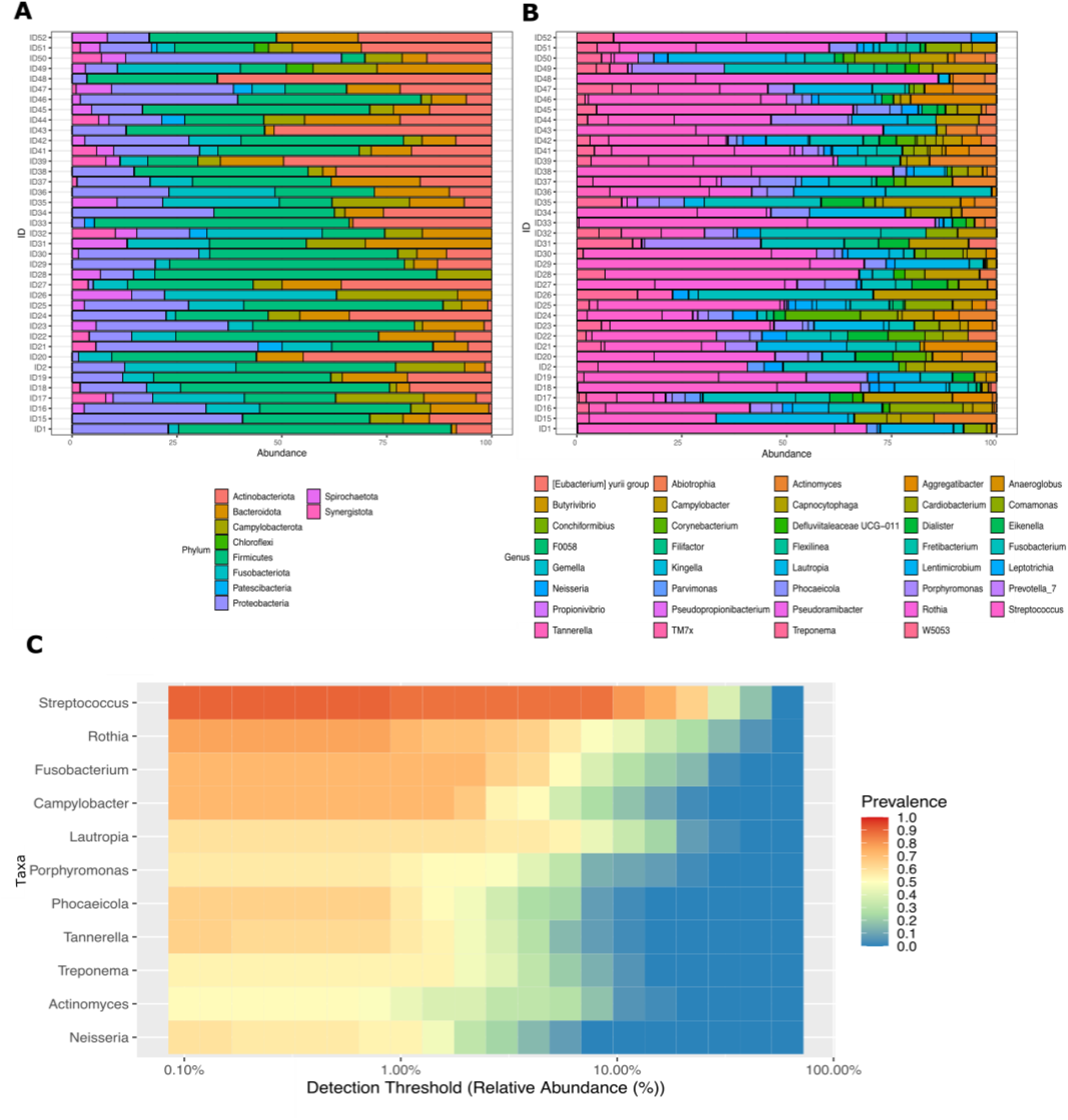
**A.** Stacked barplot showing the percentage of ASVs assigned to the phylum level over the threshold. **B.** Stacked barplot with the percentage of ASVs assigned to the genus level. **C.** Heatmap showing the ASV relative abundance on detection threshold values using a prevalence of 0.5.

### Taxonomic differences among samples are associated with different dietary habits

Intra-sample variability was compared across samples through principal coordinate analysis, showing that the microbial composition slightly changed across individuals. Four main clusters were identified through hierarchical clustering, based both on taxonomic differences and on differential abundances (Figure 3A) (C1 = 16 samples, C2 = 7 samples, C3 = 7 samples, C4 = 9 samples). The analysis of alpha-diversity revealed a high degree of similarity and higher variability of C1 and C2 compared to C3 and C4, which were also similar in composition to each other (Supplementary Figure 2). The prevalence of *Fusobacterium*, *Tannerella*, *Rothia*, *Streptococcus*, and *Lautropia* spp. significantly varied across the clusters. These genera fell into five of the main phyla composing the oral microbiome (Firmicutes, Proteobacteria, Bacteroidetes, Fusobacteria, and Actinobacteria), accounting for most of the diversity of whole microbial communities. In particular, *Fusobacterium* and *Tannerella* were both significantly enriched in C4, while *Streptococcus*, *Rothia*, and *Lautropia* were respectively enriched in C1, C2, and C3 (Figure 3B and Supplementary Figure S3).

**Figure 3.**
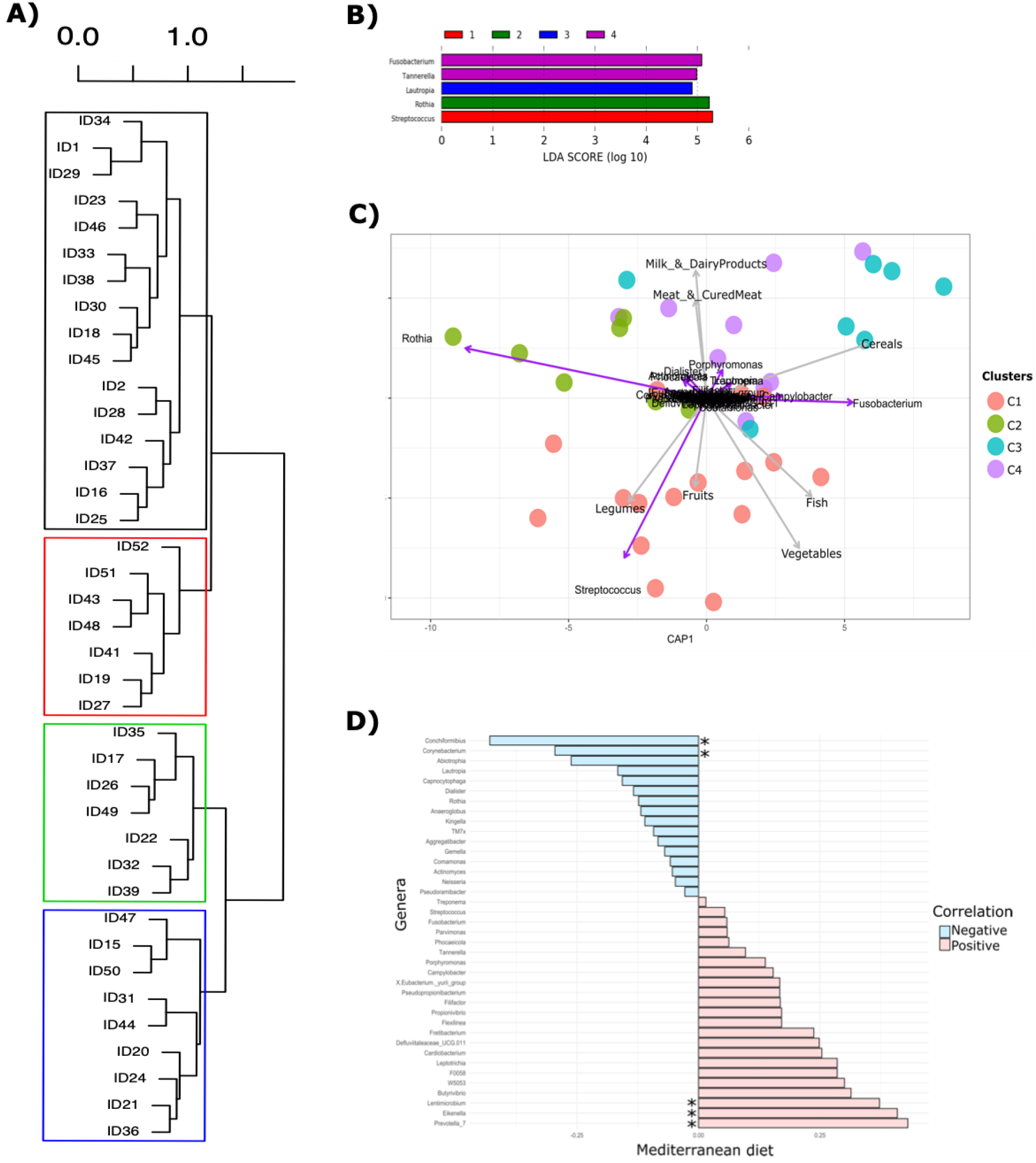
**A.** Hierarchical tree of the 39 samples based on both ASV using the Bray–Curtis distance matrix between samples and the Ward D2 criterion for computing branch disposal. Clusters were calculated by bootstrapping the gap statistic with the PAM algorithm (500 bootstraps). **B.** Linear discriminant analysis (LEfSe) results showing the score of each significant variable (genus) that better represents a given cluster. **C.** Redundancy analysis (RDA) triplot showing relationships between sample taxonomical distances divided per cluster, quantitative variables (relative abundances at the genus level), and food metadata categorical variables. **D.** Spearman correlation test for the Mediterranean score against all genus-level taxa.

In order to investigate any association between the taxonomic clusters and the collected variables, a redundancy analysis (RDA) was performed. We found that dietary variables (type of food consumed) were significantly associated with the taxonomic composition of our dataset. Specifically, legumes, fruit, vegetables, and fish were associated with C1; cereals pointed to C3; while milk, meat, and their derivatives had a stronger association with C4 (Figure 3C). Furthermore, groups related with higher consumption of fibers (vegetables, fruit) were also enriched with the *Streptococcus* genus, while C3 and C4, which were mainly associated with *Tannerella* and *Porphyromonas*, were characterized by diet rich in animal proteins (e.g., milk and meat) and higher cereal consumption. Finally, we did not detect any relevant associations between the Mediterranean score (based on the overall food consumption) and the taxonomical and clustering data (Supplementary Figure S4). However, we checked for correlations between the Mediterranean score and each bacterial genus, and we found that Mediterranean values negatively correlated with *Corynebacterium* and *Conchiformibius*, while they positively correlated with *Eikenella*, *Prevotella*, and *Lentimicrobium* (Figure 3D).

### Functional analysis and microbiota pathways

Next, we performed a cluster analysis on the KEGG profile obtained from each individual to study the microbiota activity inside the dental calculus and reveal potential associations between diet type and bacterial functions. We observed two functional clusters (FCs) according to gap statistics: FC1 was composed of 20 individuals, while FC2 included 19 individuals. Notably, FC1 was mostly composed of the taxonomy-based C1, while FC2 mainly included components belonging to C2, C3, and C4 (Figure 4A,B). The two functional clusters were differentially enriched by important pathways involved both in catabolic and anabolic processes, allowing a possible reconstruction of the different functions of microbes ascribed to the two groups. Specifically, glycan, carbohydrate, and nucleotide metabolism were enriched in FC1, while FC2 was mainly characterized by amino acid, lipid, and transport metabolism (Figure 4C). In addition, level 3 KEGG categories allowed us to gain further details on the differential enrichment of many pathways (Supplementary Figure S5). For instance, FC1 was enriched in many pathways strongly related to diet type, such as starch and sucrose metabolism, as well as amino sugars, nucleotide sugars, galactose, fructose, and mannose metabolism, which may suggest a more carbohydrate- and fiber-oriented diet, while FC2 was mostly characterized by metabolism of amino acids such as histidine, alanine, aspartate, and glutamate, which may point to the consumption of a high-protein diet, further supporting the information retrieved from the taxonomic analysis.

**Figure 4.**
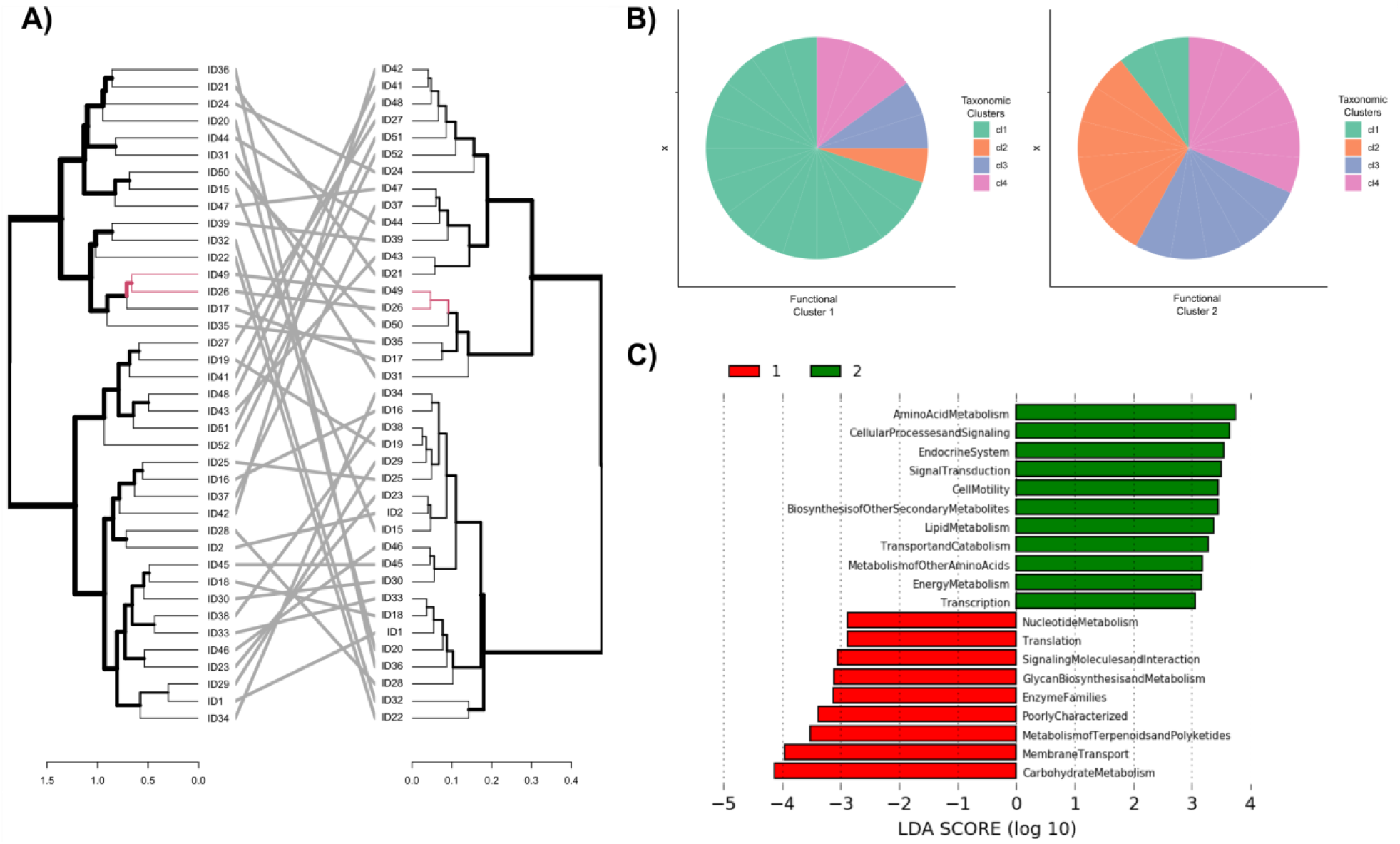
**A.** Tanglegram mixing two hierarchical cluster graphs: the left tree represents the taxonomy-based dendrogram, while the right tree was computed using distances calculated on functional profiles for each sample. **B.** Pie charts showing the inclusion partitions of each of the four taxonomy-based clusters (C1–4) inside the two functional-based clusters (FC1,2). **C.** Linear discriminant analysis using L2 KEGG ontology categories checking the significant difference between the two functional clusters.

## Discussion

The complexity of the oral microbiome is a result of the diverse microbial communities that grow in different and specific areas of the mouth, generating separate but still interconnected, biological niches (e.g., saliva, tongue, gingiva)^19^. The present study shows that the microbiome colonizing the dental calculus falls within the known variability for the oral microbiota. Indeed, the taxonomic composition comprised 10 out of the 13 phyla commonly belonging to the oral microbiome signature^29^ as well as a conspicuous number of representative genera (39). Among these, it is interesting to observe the concomitant presence of typical commensals, mainly associated with a healthy core microbiota (e.g., *Streptococcus, Fusobacterium, Rothia*, and *Actinomyces*) and of taxa associated with a higher risk of developing periodontal diseases, such as *Treponema, Tannarella*, and *Porphyromonas*, being flagged as microorganisms of the red complex^30,31^. Intriguingly, all these pathogens were found to be highly prevalent across our samples, making them relevant members of the overall core dental calculus microbiome. Still, the main genus of the microbiota was represented by *Streptococcus*, which includes both commensal and opportunistic microbes, such as *Streptococcus mutans* and the *Streptococcus anginosus* group^32,33^.

The first aim of this study was to investigate the level of inter-variability in the microbial composition of dental calculus among individuals of a rather homogeneous population. The identification of four taxonomy-based clusters was indicative of a diverse microbiota profile among samples. All the representative genera of each cluster belonged to different phyla, suggesting how phylogenetically distant microorganisms could differentially adapt to this specific ecological niche and possibly different lifestyles. The second element we wished to address in this study was whether the observed inter-variability had any association with specific dietary habits. Indeed, we found that a diet rich in viscous fibers, such as those in fruit, vegetables, and legumes, was significantly associated with C1, which was mainly represented by the *Streptococcus* genus. These specific types of fiber have been shown to reduce the postprandial increase in blood glucose and to reduce LDL cholesterol^34,35^. Furthermore, such food products are considered part of the healthy Mediterranean diet, and they are characterized by a low saturated fatty acid content, low glycemic indices^36^, and a high concentration of polyunsaturated fatty acids such as omega-3 (PUFA); furthermore, their composition includes vegetal proteins and a wide range of functional metabolites, such as polyphenols, pigments (e.g., carotenoids), and glucosinolates, with consistent nutritional values^37,38^. The high presence of *Streptococcus* among the subjects belonging to the enriched vegetable and legume consumption group (i.e., C1), might be associated with the ability of different *Streptococcus* species to express amylase-binding proteins, which would bind to host salivary α-amylases and thus exploit dietary starch for their nutrition^39–41^. It has been widely demonstrated that a high-fiber diet benefits the proliferation of many species of microorganisms, resulting in high microbial diversity^42^. Conversely, high consumption of meats, cured meats, and dairy products, which are considered signatures of a Western-based diet, is associated with higher caloric content, increased amino acid intake, and a reduction in microbial biodiversity^43^. In this study, such nutrients were mostly associated with C4 and, to a lesser extent, with C3, which were characterized by reduced biodiversity and a moderate association with some opportunistic pathogens, such as *Fusobacterium, Porphyromonas*, and *Treponema*^44,45^. Remarkably, the microbiota diversity of C1 and C2 was significantly higher compared to that of C3 and C4. These data seem to fit the cited association model of higher fiber intake with increased microbiota richness and higher meat consumption with decreased biodiversity^42,46^. As for C2, this category can be considered an admixture of the characteristics of C1 and the other groups, as it does not show signs of strong associations with a particular diet.

Interestingly, the functional analysis showed a significant association of FC1 with carbohydrate metabolism, terpenoid metabolism, and glycan pathways; thus, it is in accordance with the collected dietary information. Conversely, FC2 was characterized by enriched amino acid metabolism, energy-related pathways, and lipid metabolism. Such characteristics reflect and partially corroborate the statements made for the taxonomy-based clusters. Indeed, considering that FC1 mainly overlaps with C1, these data support the effects observed on the taxonomic level by diet: the higher carbohydrate, glycan, and terpenoid metabolism could be explained by the association with a vegetable- and fruit-rich diet, which characterized ~65% of FC1. In particular, fibers and glycans, highly present in a vegetable-based diet, are considered to play an important role in shaping oral microbiota metabolism^47^, producing an increase in all the carbohydrate processing pathways, such as fructose, galactose, and mannose metabolism, all of which were enriched in FC1. In addition, some of the positive effects driven by the Mediterranean diet in this group may be due to the activity of essential-oil-derived compounds such as terpenoids, which were found to have beneficial effects on microbiota composition, resulting in a reduced risk of oral cancer^48^. However, FC2, showing a heterogeneous sample composition comprising ~55% of both C3 and C4, reflected amino acid and fatty acid pathway enrichment. By way of a deeper assessment, regarding L3-associated pathways, it was possible to observe an enrichment in FC2 of several pro-inflammatory pathways. Indeed, a number of virulence factors, such as bacterial motility proteins, cell motility and secretion, and bacterial chemotaxis^49^, were increased within this group, along with antibiotic biosynthesis (i.e., Novobiocin), xenobiotics metabolism^50^, and glutathione metabolism^51^, which help in adaptation to stress. Other signals of possible dietary connection with the dental calculus microbiome composition came from the correlation of the value of adherence to the Mediterranean diet with some of the genera found in our cohort. Particularly, we suggested that *Prevotella, Eikenella*, and *Lentimicrobium* seem to be positively associated with greater adherence to a Mediterranean diet. This result is in accordance with what has already been observed in other studies in the positive selection exerted by high-fiber and -vegetable diets on *Prevotella* spp. and *Eikenella*^52,53^.

In conclusion, the evidence obtained in the present pilot study suggests that the dental calculus microbiome accounts for a very biodiverse environment. In particular, this microbiome is representative of the ecological niche that constitutes the oral cavity, and, moreover, its composition and gene contents may be driven by specific dietary elements. This is, to our knowledge, the first evidence of an association between the microbiome composition of dental calculus and food consumption, suggesting its use as a possible signature of individual nutrition, probably even on ancient populations. Further investigation is needed to compare the effects of different dietary regimens (such as vegan or vegetarian) and health status in order to highlight other types of individual information that could leave a trace (i.e., biomarker) on the dental calculus microbiome.

## Acknowledgments

We thank the BMR Genomics srl (Padova, Italy), and Dr. E.Sattin for their support in DNA sequencing.

