## Supplementary Tables for "Assessing the influence of the Mediterranean diet on dental calculus microbiome composition: a pilot study"

| | Fruit # | Vegetables # | Legumes * | Cereals # | Fish * | Red meat and Salumes * | Milk and dairy products # | Alcohol# | Olive oil |
|---|---|---|---|---|---|---|---|---|---|
| ID1 | >2 | >2 | <1 | <1 | >2 | <1 | <1 | <1 | regularly |
| ID15 | <1 | 1_2 | 1_2 | >2 | 1_2 | 1_2 | 1_2 | >2 | regularly |
| ID16 | 1_2 | 1_2 | 1_2 | 1_2 | 1_2 | 1_2 | 1_2 | <1 | regularly |
| ID17 | 1_2 | 1_2 | 1_2 | >2 | 1_2 | 1_2 | >2 | <1 | regularly |
| ID18 | 1_2 | >2 | 1_2 | 1_2 | 1_2 | >2 | <1 | <1 | regularly |
| ID19 | 1_2 | 1_2 | 1_2 | 1_2 | 1_2 | 1_2 | 1_2 | 1_2 | often |
| ID2 | <1 | 1_2 | >2 | >2 | 1_2 | <1 | <1 | 1_2 | regularly |
| ID20 | <1 | <1 | 1_2 | 1_2 | 1_2 | 1_2 | 1_2 | <1 | regularly |
| ID21 | 1_2 | >2 | 1_2 | 1_2 | 1_2 | 1_2 | 1_2 | 1_2 | regularly |
| ID22 | 1_2 | 1_2 | <1 | <1 | >2 | 1_2 | 1_2 | <1 | regularly |
| ID23 | 1_2 | 1_2 | <1 | >2 | >2 | 1_2 | <1 | <1 | often |
| ID24 | 1_2 | 1_2 | 1_2 | 1_2 | 1_2 | 1_2 | <1 | 1_2 | regularly |
| ID25 | 1_2 | >2 | 1_2 | 1_2 | >2 | >2 | <1 | 1_2 | regularly |
| ID26 | <1 | 1_2 | <1 | >2 | 1_2 | 1_2 | <1 | <1 | regularly |
| ID27 | 1_2 | 1_2 | 1_2 | 1_2 | 1_2 | >2 | <1 | <1 | regularly |
| ID28 | 1_2 | >2 | >2 | 1_2 | 1_2 | <1 | <1 | <1 | regularly |
| ID29 | 1_2 | 1_2 | 1_2 | 1_2 | 1_2 | 1_2 | 1_2 | <1 | regularly |
| ID30 | <1 | 1_2 | 1_2 | 1_2 | <1 | 1_2 | >2 | <1 | often |
| ID31 | 1_2 | 1_2 | <1 | 1_2 | 1_2 | >2 | 1_2 | 1_2 | regularly |
| ID32 | 1_2 | >2 | 1_2 | 1_2 | >2 | 1_2 | 1_2 | <1 | regularly |
| ID33 | 1_2 | 1_2 | 1_2 | 1_2 | 1_2 | 1_2 | <1 | 1_2 | often |
| ID34 | 1_2 | 1_2 | 1_2 | 1_2 | 1_2 | 1_2 | 1_2 | 1_2 | regularly |
| ID35 | 1_2 | 1_2 | 1_2 | 1_2 | 1_2 | 1_2 | 1_2 | <1 | occasionally |
| ID36 | <1 | 1_2 | 1_2 | 1_2 | 1_2 | 1_2 | 1_2 | 1_2 | often |
| ID37 | 1_2 | 1_2 | 1_2 | >2 | 1_2 | 1_2 | 1_2 | <1 | regularly |
| ID38 | 1_2 | 1_2 | >2 | 1_2 | 1_2 | 1_2 | 1_2 | <1 | regularly |
| ID39 | 1_2 | 1_2 | >2 | 1_2 | 1_2 | 1_2 | 1_2 | 1_2 | often |
| ID41 | 1_2 | 1_2 | 1_2 | 1_2 | >2 | 1_2 | 1_2 | 1_2 | regularly |
| ID42 | <1 | 1_2 | <1 | 1_2 | 1_2 | >2 | <1 | 1_2 | occasionally |
| ID43 | <1 | 1_2 | 1_2 | 1_2 | <1 | 1_2 | 1_2 | <1 | occasionally |
| ID44 | 1_2 | 1_2 | 1_2 | 1_2 | <1 | 1_2 | 1_2 | 1_2 | often |
| ID45 | 1_2 | 1_2 | 1_2 | >2 | <1 | >2 | >2 | <1 | regularly |
| ID46 | <1 | 1_2 | 1_2 | 1_2 | >2 | 1_2 | <1 | 1_2 | often |
| ID47 | 1_2 | 1_2 | <1 | <1 | 1_2 | 1_2 | 1_2 | <1 | often |
| ID48 | <1 | 1_2 | <1 | 1_2 | 1_2 | 1_2 | 1_2 | 1_2 | often |
| ID49 | 1_2 | >2 | 1_2 | >2 | 1_2 | 1_2 | 1_2 | <1 | regularly |
| ID50 | <1 | 1_2 | <1 | 1_2 | 1_2 | 1_2 | 1_2 | <1 | often |
| ID51 | 1_2 | 1_2 | 1_2 | 1_2 | 1_2 | 1_2 | 1_2 | 1_2 | often |
| ID52 | 1_2 | 1_2 | 1_2 | 1_2 | 1_2 | 1_2 | 1_2 | 1_2 | regularly |

**Table S1.** Dietary servings values for each food obtained from individual surveys. These values represent the number of servings per day or per week. Main meals include breakfast, lunch and dinner.

### The values represent the number of serving per day.

* The values represent the number of serving per week.

| | Raw reads | Post-quality trimming reads |
|---|---|---|
| **ID1** | 59218 | 46123 |
| **ID2** | 75842 | 45021 |
| **ID52** | 45039 | 26215 |
| **ID15** | 45471 | 30328 |
| **ID16** | 61455 | 36614 |
| **ID17** | 116924 | 72566 |
| **ID18** | 86294 | 56820 |
| **ID19** | 66675 | 42483 |
| **ID20** | 31494 | 19878 |
| **ID21** | 62031 | 39079 |
| **ID22** | 76161 | 34325 |
| **ID23** | 65128 | 42160 |
| **ID24** | 43842 | 25685 |
| **ID25** | 108435 | 67620 |
| **ID26** | 88112 | 50076 |
| **ID27** | 59341 | 36215 |
| **ID28** | 49139 | 30146 |
| **ID29** | 70877 | 50128 |
| **ID30** | 69953 | 46332 |
| **ID31** | 78873 | 40165 |
| **ID32** | 51613 | 28263 |
| **ID33** | 85634 | 57787 |
| **ID34** | 62985 | 41133 |
| **ID35** | 73380 | 40934 |
| **ID36** | 35768 | 18324 |

| | | |
|---|---|---|
| **ID37** | 81853 | 46902 |
| **ID38** | 103472 | 73810 |
| **ID39** | 64373 | 35958 |
| **ID40** | 1790 | 975 |
| **ID41** | 109322 | 72009 |
| **ID42** | 88848 | 56225 |
| **ID43** | 73255 | 49121 |
| **ID44** | 53906 | 31052 |
| **ID45** | 88329 | 52591 |
| **ID46** | 71470 | 45098 |
| **ID47** | 64671 | 40188 |
| **ID48** | 46930 | 32470 |
| **ID49** | 119449 | 72398 |
| **ID50** | 81989 | 54147 |
| **ID51** | 74526 | 45584 |

**Table S2.** Summary of per-sample raw reads count, before and after quality trimming.
