## Supplementary Figures for "Assessing the influence of the Mediterranean diet on dental calculus microbiome composition: a pilot study"

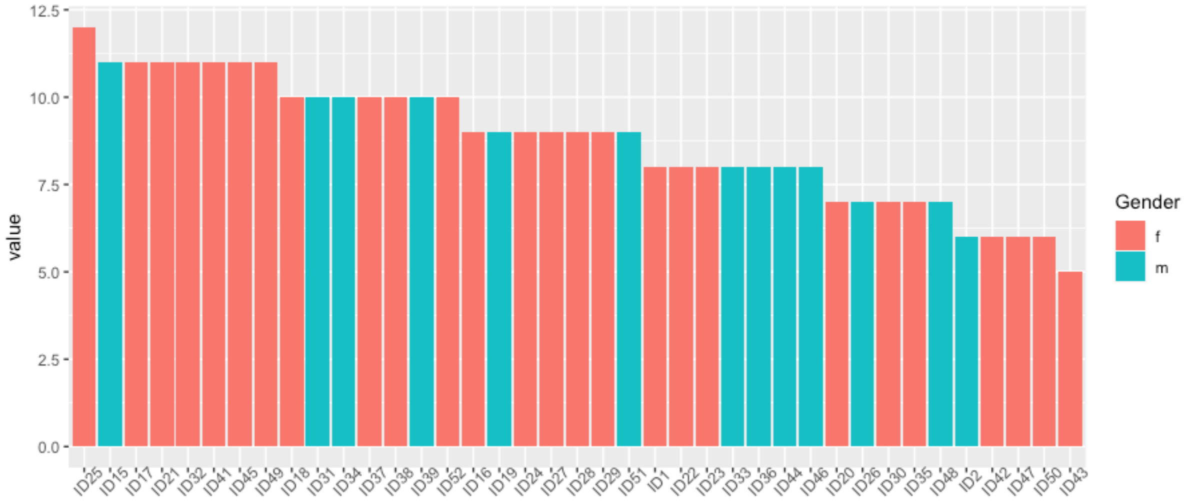


**Figure S1.** Mediterranean score for each sample, according to the KIDMED index system. The barplot also shows the gender division.

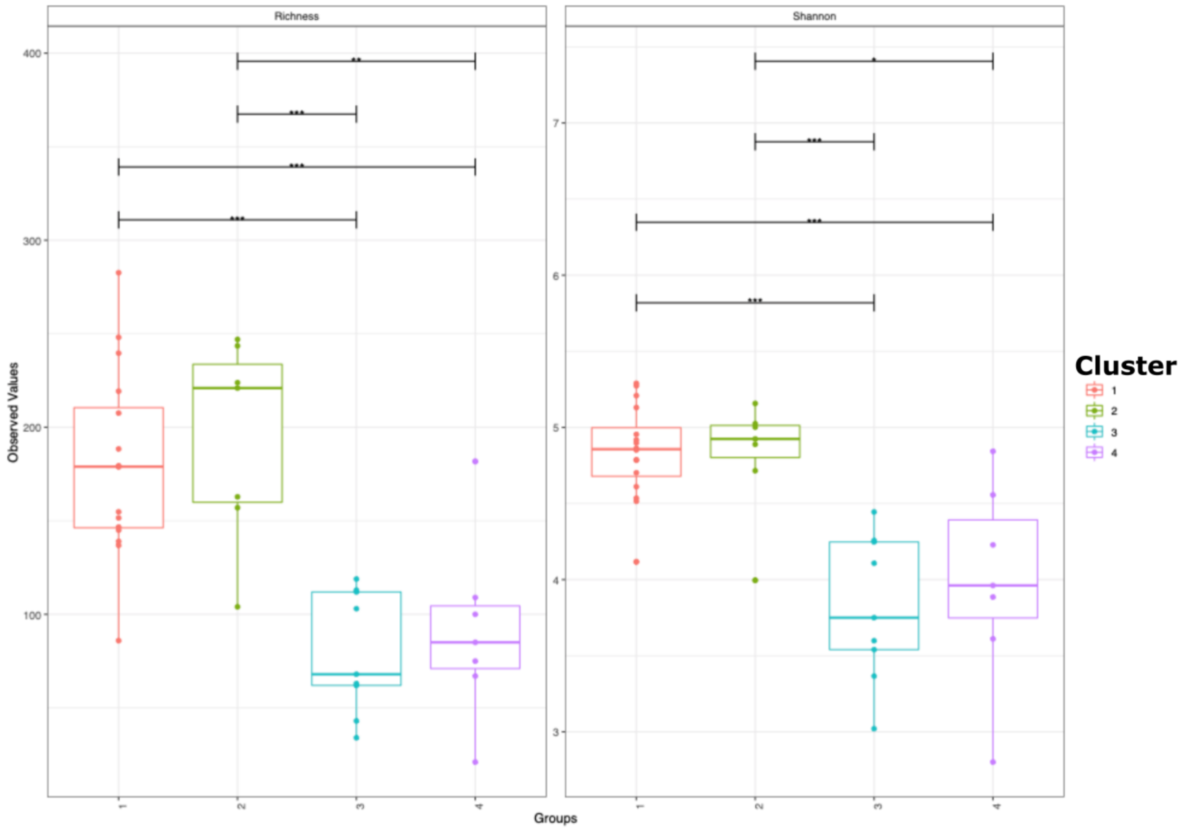


**Figure S2.** Boxplot showing the alpha diversity of each cluster using both richness and shannon indexes. Groups that are significantly different are connected by the horizontal bar.

* p < 0.05

** p < 0.01

***p < 0.001

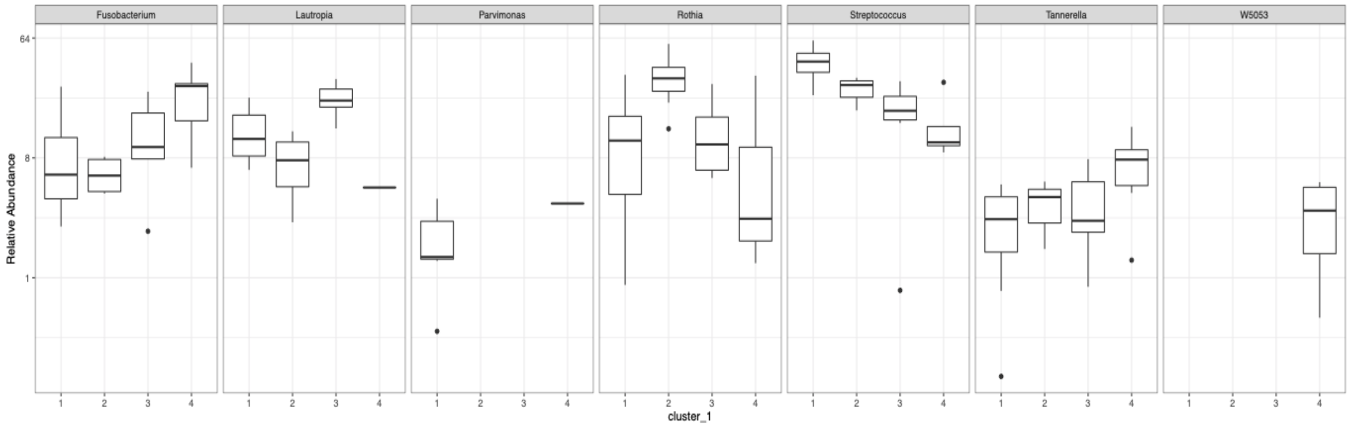


**Figure S3.** The plot shows the distribution of relative abundances of those genera that resulted significantly different in a Kruskal-Wallis test ($p < 0.01$), computed considering the clustering division (C1, C2, C3, C4). *Parvimonas* and *Peptostreptococcaceae bacterium oral taxon 113* are only present in high abundance only in C1 and C4, so they were removed in the LEfSe analysis shown in Figure 3B in the paper.

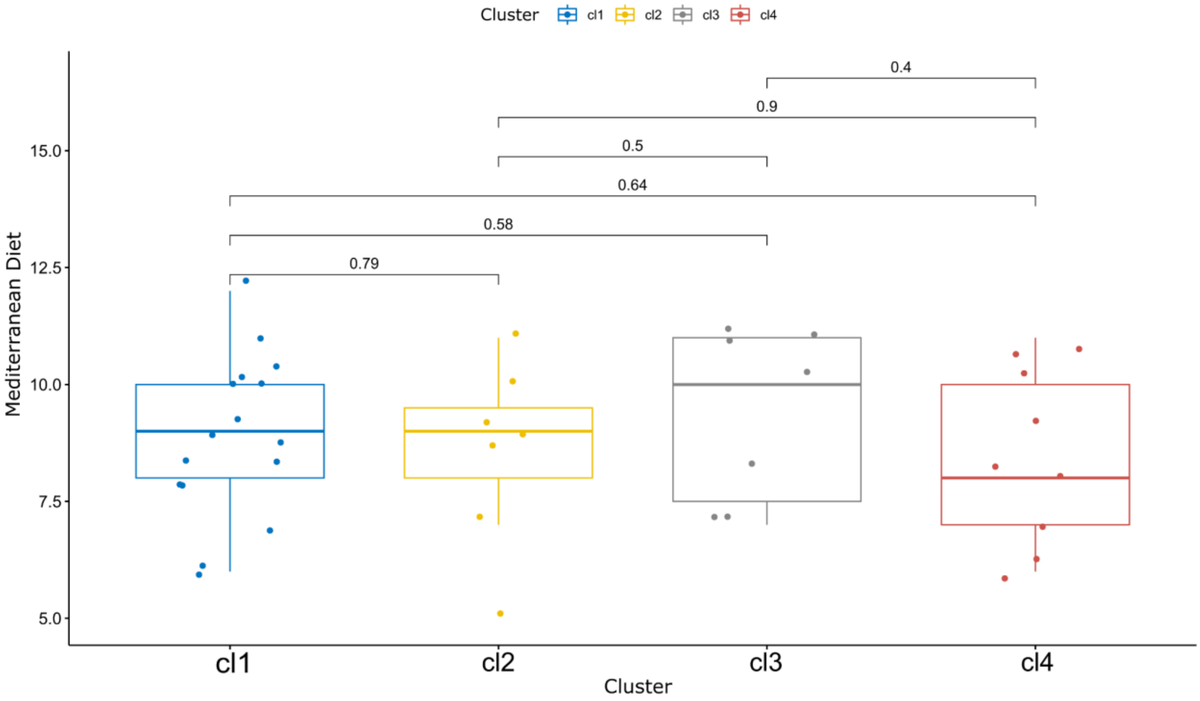


**Figure S4.** Differences in distribution of Mediterranean score values along the clustering division (cl). The significance was computed using a t-test.

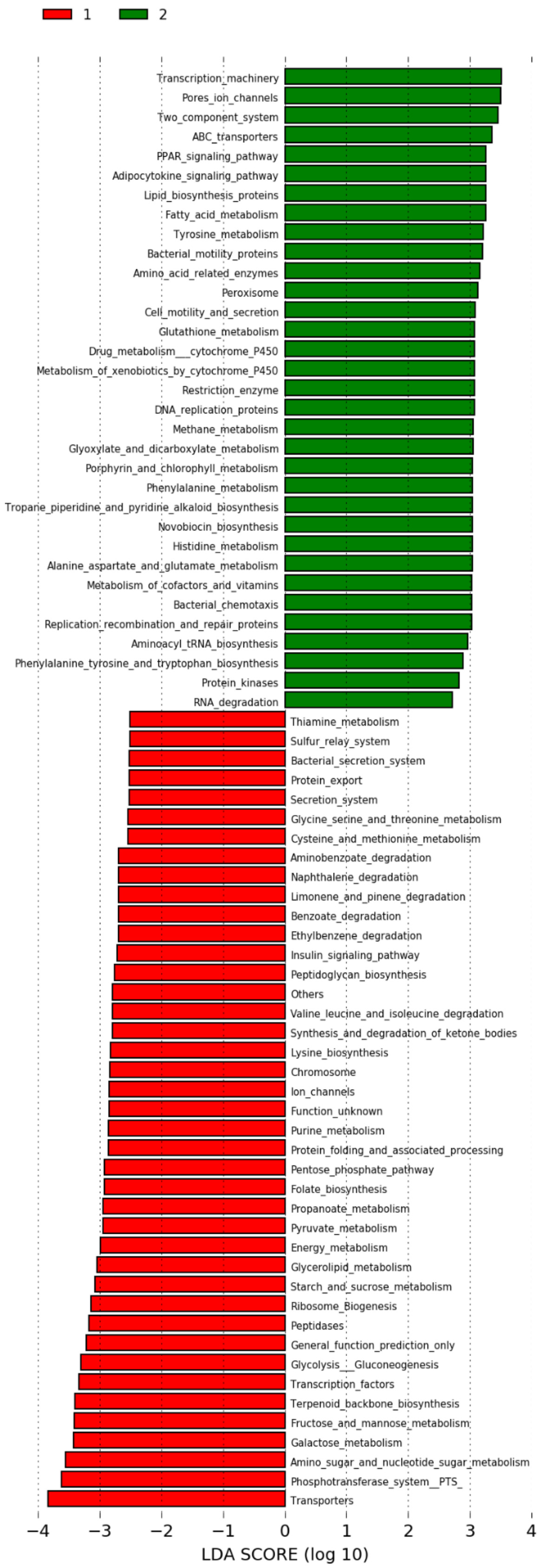


**Figure S5.** Result of Linear Discrimination Analysis on FC1 (green) vs FC2 (red) using L3 KEGG ontology database annotations.
